# Impact of soil salinity on the cowpea nodule-microbiome and the isolation of halotolerant PGPR strains to promote plant growth under salinity stress

**DOI:** 10.1101/856765

**Authors:** Salma Mukhtar, Ann M. Hirsch, Noor Khan, Kauser A. Malik, Ethan A. Humm, Matteo Pellegrini, Baochen Shi, Leah Briscoe, Marcel Huntemann, Alicia Clum, Brian Foster, Bryce Foster, Simon Roux, Krishnaveni Palaniappan, Neha Varghese, Supratim Mukherjee, T.B.K. Reddy, Chris Daum, Alex Copeland, Natalia N. Ivanova, Nikos C. Kyrpides, Nicole Shapiro, Emiley A. Eloe-Fadrosh, Maskit Maymon, Muhammad S. Mirza, Samina Mehnaz

## Abstract

Four soil samples (SS-1—SS-4) isolated from semi-arid soils in Punjab, Pakistan were used as inocula for cowpea (V*igna unguiculata* L.) grown under salinity stress to analyze the composition of bacteria in the rhizosphere and within nodules through cultivation-dependent and cultivation-independent methods. Two cowpea varieties, 603 and the salt-tolerant CB 46, were each inoculated with four different native soil samples, and data showed that plants inoculated with soil samples SS-2 and SS-4 grew better than plants inoculated with soil samples SS-1 and SS-3. Bacteria were isolated from both soils and nodules, and 34 of the 51 original isolates tested positive for PGPR traits in plate assays with many exhibiting multiple plant growth-promoting properties. A number of isolates were positive for all PGPR traits tested. For the microbiome studies, environmental DNA (eDNA) was isolated from SS-1 and SS-4, which represented the extremes of the Pakistan soils to which the plants responded, and by 16S rRNA gene sequencing analysis were found to consist mainly of Actinobacteria, Firmicutes, and Proteobacteria. However, sequencing analysis of eDNA isolated from cowpea nodules established by the trap plants grown in the four Pakistan soils indicated that the nodule microbiome consisted almost exclusively of Proteobacterial sequences, particularly *Bradyrhizobium*. Yet, many other bacteria including *Rhizobium*, *Mesorhizobium, Pseudomonas,* as well as *Paenibacillus*, *Bacillu*s as well as non-proteobacterial genera were isolated from the nodules of soil-inoculated cowpea plants. This discrepancy between the bacteria isolated from cowpea nodules (Proteobacteria and non-Proteobacteria) versus those detected in the nodule microbiome (Proteobacteria) needs further study.

## Introduction

The cowpea (*Vigna unguiculata* L.) cultivar Lobia is one of the most important legumes cultivated in semi-arid regions of Pakistan. It is produced and consumed by subsistence farmers, particularly in the barani (dry-farming) zones in Pakistan. A multipurpose crop, cowpea is used as a pulse, forage, and vegetable crop (Muhammad et al. 1993; Imran et al. 2012), and also plays an important role in crop rotation systems and restoration of soil fertility through nitrogen fixation. Environmental factors—air temperature, soil salinity, moisture content, soil fertility, and photoperiod length—all have a significant influence on growth and yield of cowpea (Ali et al. 2004). To ensure that cowpea remains a key crop as the climate changes and potentially disrupts dryland farming, the agriculture department of Punjab, Pakistan is importing exotic cowpea varieties and running plant trials under local conditions in indigenous soils.

Rhizobia are soil-inhabiting Gram-negative bacteria (Proteobacteria) that initiate root nodule development on legume plants where they are maintained as intracellular symbionts within the nodules. There they convert atmospheric nitrogen into ammonia for assimilation by the plant in exchange for plant-derived organic acids (Garg and Geetanjali 2007; Graham, 2008; Sulieman and Tran 2014). The majority of bacteria that fix nitrogen in root nodules of leguminous plants are members of the α-Proteobacteria, *Rhizobium, Bradyrhizobium, Ensifer*, *Phyllobacterium*, *Mesorhizobium*, *Devosia*, *Azorhizobium*, *Allorhizobium* (Hirsch 1992; Sawada et al. 2003; Lafay and Burdon 2007) and also *Microvirga* (Ardley et al. 2012). Among the β-Proteobacteria, members of the *Burkholderiaceae*, known as beta-rhizobia, also nodulate legumes (Bontemps et al. 2010, Gyaneshwar et al. 2011, Estrada de los Santos et al. 2018).

A vast literature on microbes other than rhizobia detected within nitrogen-fixing nodules has accumulated, starting with the earliest report made by Beijerinck (1888) who described the isolation of brown and yellow-colored *Bacillus-*like strains from root nodules. Since then, culture-dependent studies have pointed to a much higher diversity of bacteria associated with root nodules beyond rhizobia. Many of these bacteria do not fix nitrogen and are not members of the Proteobacteria (Deng et al. 2011; Aserse et al. 2013). These bacteria have been called non-rhizobial endophytes (NRE) or nodule-associated bacteria (NAB) (Velázquez et al. 2013), and include *Bacillus, Enterobacter, Agrobacterium, Acinetobacter, Paenibacillus, Pantoea, Mycobacterium, Micromonospora*, and *Pseudomonas* (Martínez-Hidalgo and Hirsch 2017 and references therein). They are not involved in nodule formation *per se*, but many have plant growth-promoting abilities including phytohormone production or mineral solubilization activity (P, Zn) either by themselves or when co-inoculated with rhizobia and other bacteria (Schwartz et al. 2013; Gupta et al. 2015). Most of the NRE are not pathogenic, but some may be serious pathogens of animals or plants, e.g., *Staphylococcus, Bordetella* (Sturz et al. 1997; Xu et al. 2014) and species of *Burkholderiaceae* such as *B. cepacia* (Eberl and Vandamme 2016). In the last few years, many of the symbiotic *Burkholderia* species (Gyaneshwar et al. 2011) have been placed into new genera, e.g., *Paraburkholderia* and *Trinickia* (Sawana et al. 2014; Estrada de los Santos et al. 2018).

Although semi-arid regions in Pakistan have a large biodiversity of rhizobial resources, only a few of them have been extensively studied using molecular approaches and thus the microbial diversity of cowpea-nodulating rhizobia and associated nodule bacteria in semi-arid soils remains largely unexplored. Our goal was to assess the effect of native soil microbial communities on the composition of the cowpea nodule microbiome under various salinity conditions. We predicted that we would find bacteria with plant growth-promoting characteristics in addition to rhizobia within cowpea root nodules inoculated with soil from Pakistan, and that these arid-soil adapted strains could enhance the rhizobia-legume interaction under climate-change conditions.

## Materials and methods

### Soil sampling and physicochemical analysis

Mandi Bahauddin, a fertile agricultural belt where the main crops are wheat, maize, sugarcane, chickpea, cowpea, and tobacco, is a district of Punjab, Pakistan. It is situated at 32°35′10″ N, 73°36′12″ E and is bordered in the northwest by the Jhelum river and in the southeast by the Chenab River (Fig. S1). Cowpea fields that covered an area of approximately 1.5 km made up the study site and the sampling areas were selected according to land use and vegetation cover. At each site, approximately 1 kg from different soils (SS-1—SS-4) collected at a 15-cm depth from four different locations was placed into black sterile polyethylene bags. Rhizosphere soil samples were collected by gently removing the plants and obtaining the soil attached to the roots.

Soil physical and chemical properties were determined for each sample based on 400 g of dried and sieved soil. Electrical conductivity (dS/m) was measured by using 1:1 (w/v) soil to water mixtures at 25°C (Adviento-Borbe et al. 2006). The pH was determined by using 1:2.5 (w/v) soil to water suspension. Moisture (%), temperature (°C), and texture class were assessed by the methodology of Anderson et al. (1993), and organic matter (OM) concentration (C_org_) was calculated by the Walkley-Black method (Walkey and Black 1934). Phosphorous was estimated by extraction with sodium bicarbonate (Olsen et al. 1954), and calcium and magnesium were detected by atomic absorption spectrometry. Nitrate ions were measured by the Raney-Kjeldahl method and potential acidity (H^+^Al) was determined by an equation based on the pH in SMP buffer solution (pH SMP). Cation exchange capacity (CEC) is the capacity to retain and release cations (Ca^2+^, Mg^2+^, K^+^ and Na^+^) and the sodium adsorption ratio (SAR) is the measure of soil sodicity, which is calculated as the ratio of sodium to magnesium and calcium.

### Soil eDNA extraction

Environmental DNA (eDNA) was isolated from two of the four rhizosphere soil samples, SS-1 and SS-4, using the QIAGEN DNeasy Power Soil Pro DNA isolation kit. Two PowerBead tubes containing 0.5 g for each sample were used for the isolation. The eDNA was sent to MR DNA (Shallowater, TX) for sequencing using the Illumina platform (San Diego, CA).

### Soil-inoculated plant trap experiment

Cowpea plants (603; PI 339603 and CB 46; PI 548784) were grown in plastic pots containing 300 g of a 2:1 vermiculite:perlite mixture with 5 g of cowpea rhizospheric soil added to it. This was then topped with sterilized polyethylene beads to minimize contamination. Cowpea 603 is a slightly salt-tolerant variety whereas cowpea CB 46 is highly salt-tolerant (Helms et al. 1991). The first set of pot experiments was performed with cowpea 603 seeds, which were inoculated with four different soil samples under normal soil conditions. In the second and third set of experiments, cowpea 603 and CB 46 seeds were inoculated with four different soil samples and grown under slight salinity stress conditions (ca. 1% NaCl) under controlled greenhouse conditions at 21-30°C during the day and 17-22°C at night in the UCLA Plant Growth Center. Inoculated plants were watered with ¼-strength Hoagland solution without nitrogen whereas control plants were watered with ¼-strength Hoagland solution with or without nitrogen or with sterile water. Three pots with 6 plants/pot were maintained for the control, and 6 pots with 6 plants each for the treatment sets. All plants were harvested after 8 weeks, and data for plant growth parameters such as biomass, shoot length, and nodule number per plant were recorded and statistically analyzed using the Statistical Package for the Social Sciences (SPSS) software (IBM Statistics 23.0).

### Isolation of bacteria from root nodules of cowpea for cultivation-dependent and cultivation-independent analyses

Root nodules were surface sterilized in 10% bleach for 5 min, washed with sterile water 10 times, and then squashed in 50 μL of sterile water with a sterile glass rod. The nodule squashate was serially diluted from 10^−1^ to 10^−5^. Approximately 50 μL of dilutions ranging from 10^−3^ to 10^−5^ were spread onto 3 different culture media: TY (Tryptone Yeast Extract), TSA (Tryptone Soya), and AG (Arginine Glycerol) agar, and plates incubated at 30°C usually for 2-3 days. However, for selecting slow growing bacteria like the actinomycetes, the plates were incubated for about a week. Bacterial colonies were counted and number of bacteria per gram of sample was calculated. The bacterial isolates were repeatedly sub-cultured to obtain single colonies, which were grown in TY, TSA, and AG broth and stored in 15% glycerol at −80°C for subsequent characterization.

For the cultivation-independent analysis, surface-sterilized nodules from plants grown in the different saline soils were crushed as described above, and eDNA was extracted from the crushed nodules using the QIAGEN PowerSoil DNA Isolation Kit. The eDNA was stored at −20°C and sent to the Joint Genome Institute for paired-end sequencing using the KAPA-Illumina library creation kit (KAPA Biosystems) and the HiSeq-2500 Illumina platform (San Diego, CA). A data cleaning process was applied to all sequences prior to analysis. Low-quality bases with a Phred quality value lower than 20 were trimmed off the read ends. To determine the taxonomic composition of the microbiome, the 16S rRNA gene sequences were extracted from the metagenomic shotgun sequencing data using a mapping-based method similar to that described by Shi et al. (2015). The sequences were aligned with paired reads against the rRNA database (Greengenes v13.5, non-redundant precalculated OTU references, 97_otus from PICRUSt) (DeSantis et al. 2006; Langille et al. 2013). The alignments were performed using Bowtie2 (Langmead et al. 2012) to identify sequences with the best hits having ≥80% nucleotide identity to the references. The annotation of the extracted 16S rRNA gene sequences was refined by cross-referencing the NCBI Reference Sequence (Pruitt et al. 2005). The microbial relative abundances in each sample were calculated on the basis of the 16S rRNA gene sequences of the classified taxa.

### Morphological and physiological characterization of bacterial isolates

Bacterial colonies were characterized on the basis of color, shape, size, margin, and elevation. The bacterial cell shape and motility determinations were done using light microscopy. Bacterial isolates were grown in the presence of different salt concentrations (0-5% NaCl) at pH ranges of 6-10 and temperatures of 4-42°C to assess their abiotic stress tolerance potential.

### Amplification, purification, and sequencing of the 16S rRNA Gene

Bacteria were suspended from a single colony of the selected isolates grown on TY plates into 20 μl of sterile-filtered water. The 16S rRNA gene was amplified by PCR using the forward primer fD1 and the reverse primer rD1 (Weisburg et al. 1991). Amplification was performed in a total volume of 20 μL containing 15 μL sterile-filtered water, 2 μL of bacterial sample, 2 μL of 10× Taq Buffer (MgCl_2_), 0.2 μL fD1 and rD1 (3.2 pmole/μL), 0.2 μL dNTPs, and 0.2 μL Taq DNA polymerase. Amplified 16S rDNA products in a 0.8% low-melting point agarose gel (100 V, 400 mA, 1 h) were visualized with ethidium bromide and purified. The gel extraction was performed with the Invitrogen Quick Gel Extraction Kit according to the manufacturer’s directions. Purified PCR products were commercially sequenced by using forward and reverse primers (Laragen, USA).

### Phylogenetic analysis

Acquired sequences were assembled and analyzed with the help of Chromus Lite 2.01 sequence analysis software (Technelysium Pty Ltd. Australia). The gene sequences were compared to those deposited in the GenBank nucleotide database using NCBI BLAST. Sequences were aligned using Clustal X 2.1 and a phylogenetic tree was constructed using the neighbour-joining method (Saitou and Nei 1987). Bootstrap confidence analysis was performed on 1000 replicates to determine the reliability of the distance tree topologies obtained. The evolutionary distances were computed using the Maximum Composite Likelihood method (Tamura et al. 2004) and are presented in the units of number of base substitutions per site. All positions containing gaps and missing data were eliminated from the dataset (complete deletion option). Phylogenetic analyses were conducted in MEGA7 (Kumar et al. 2016).

## Assays for plant growth promotion

### Phosphate solubilization assay

Pikovskaya phosphate medium (PVK) was made according to Pikovskaya (1948). For inoculation onto plates, the bacterial strains were grown in liquid medium until stationary phase, followed by harvesting of cells by centrifugation (8000 × g, 10 min). The pellets were then washed three times with sterile water for removing any traces of the medium. After diluting the bacterial suspension with sterile water (OD_600_ = 0.2), ten μL of it was spotted onto the plates. The plates were then incubated at 30 °C for 3 days and the size of the clearing zone around the colony was measured.

A quantitative analysis of P-solubilization of bacterial isolates was done by the molybdate blue color method (Watanabe and Olsen 1965). Available P was calculated after 7 and 14 days. Cell-free supernatants were used for the quantification of P-solubilization. After recording the pH of the cell-free supernatants, they were filtered through 0.2 μm sterile filters (Orange Scientific GyroDisc CA-PC, Belgium) to remove any residues. Solubilized phosphates (primary and secondary orthophosphate) were measured by spectrophotometry (Camspec M350-Double Beam UV-Visible Spectrophotometer, UK) at 882 nm, and values were calculated using a standard curve (2, 4, 6, 8, 10, 12 ppm KH_2_PO_4_ solutions).

### Siderophore assay

CAS agar medium devoid of nutrients was used as an indicator of siderophore presence. The components needed for a liter of the overlay medium were as described in Perez-Miranda et al. (2007). Ten mL of a 0.9% CAS agar gel was spread as an overlay on culture plates of selected strains grown for 4 days on two solid media (LB and TY). After a maximum period of 15 min, a color change in the blue medium was observed around the colonies. The experiment was repeated three times.

### Hydrogen cyanide production assay

HCN detection was done using filter papers (soaked in 0.5% picric acid in 2% sodium carbonate), which were placed on the lid of the Petri plate (Sadasivam and Manickam 1992).

### Antifungal activity assay

The antifungal activity of the bacterial isolates was observed using a dual-culture method where the test bacteria and the fungal culture are co-incubated on a potato dextrose agar (PDA; Sigma, USA) plate (Khan et al. 2018). The antagonistic effectiveness of the test bacteria is assessed by measuring the inhibition of fungal mycelia around the bacteria. The observations were recorded when fungus attained full growth on the control plate. The fungal strains used to evaluate anti-fungal activity of the bacterial isolates were: *Fusarium oxysporum, Fusarium solani, Alternaria solani, Aspergillus flavus,* and *Curvularia* sp.

### Enzyme assays for bacterial isolates

Protease activity was tested on the medium described by Kumar et al. (2009). Amylase activity was identified by using a starch hydrolysis test (Sigmon 2008). Cellulase activity was tested by using LB and TY media containing 1% carboxy methyl cellulose (CMC), and indicated by formation of a clear zone after staining with Gram’s iodine for 3-5 minutes (Kasana et al. 2008). Lipase activity was examined by using LB and TY media with 1% butyrin and Tween 80 in a hydrolysis assay as described by Sierra (1957). Chitinase activity was detected with colloidal chitin medium as described by Kuddus and Ahmad (2013). Clear zones around the bacterial colonies are considered as positive results for chitinase (Kaur et al. 2012).

## Results

### Physicochemical properties of cowpea-planted rhizosphere soil samples collected in Mandi Bahauddin

The four soil samples were characterized by a great spatial variability and covered a significant variation in electrical conductivity (EC), soil pH, salinity, organic matter (OM), texture class, CEC (cation exchange capacity), SAR (sodium adsorption ratio) and other factors. Soil samples (SS) SS-1 and SS-3 were more saline (5.05 and 5.29 dS/m) compared to SS-2 and SS-4 (4.85 and 4.56 dS/m). SS-2 had more OM (21.23 to 31.75 g/Kg) compared to SS-3, which was more alkaline (pH 8.11) compared to the other samples (Table S1). Soil temperatures ranged from 29.2 to 38.1°C, with the highest values detected in SS-3 and the lowest in SS-1 (Table S1). Moisture ranged from 18-24% with the highest in SS-4 and the lowest in SS-1 (Table S1). The available P, K, Ca, and Mg contents were lower in SS-3 compared to SS-2 and SS-4. CEC values ranged from 58.98 to 67.52 mg/dm^3^ and SAR (Sodium Adsorption Ratio) values from 10.41 to 12.42 with the highest value in SS-1 and the lowest value in SS-2 (Table S1).

### Results of trap experiments

Trap experiments with cowpea were pursued because we hypothesized that this important crop legume would form root nodules that contained not only nitrogen-fixing bacteria, but also “helper” bacteria that promoted cowpea growth in these soils based on previous studies (Martínez-Hidalgo and Hirsch 2017). Plant growth-promoting responses such as changes in shoot length, plant biomass, and number of nodules per plant were recorded for control cowpea and for cowpea plants grown in pots under different salt stress conditions. The largest increase in shoot length occurred in cowpea 603 plants inoculated with SS-4 (Fig. 1, Fig. S2). SS-1 and SS-3-inoculated plants overlapped in shoot length with the uninoculated controls whereas both SS-2 and SS-4-grown plants exhibited shoot lengths equivalent to the +N control. Also, shoot length increased slightly more in control soil-inoculated cowpea 603 plants than in cowpea 603 and CB 46 plants grown under salinity stress (Fig. 1). Cowpea 603 plants grown in +N medium without salt were green and healthy whereas when they were grown in SS-1 and SS-3, the 603 plants were yellow and showed limited root development, most likely because of the higher salinity of these soils. The salt-tolerant CB 46 cowpea was less affected by the SS-1 and SS-3 soils than cowpea 603, and in SS-2 and SS-4 soils, both cultivars were more similar in appearance to the no-salt control even when grown under salt stress (Fig. S2).

**Fig. 1.**
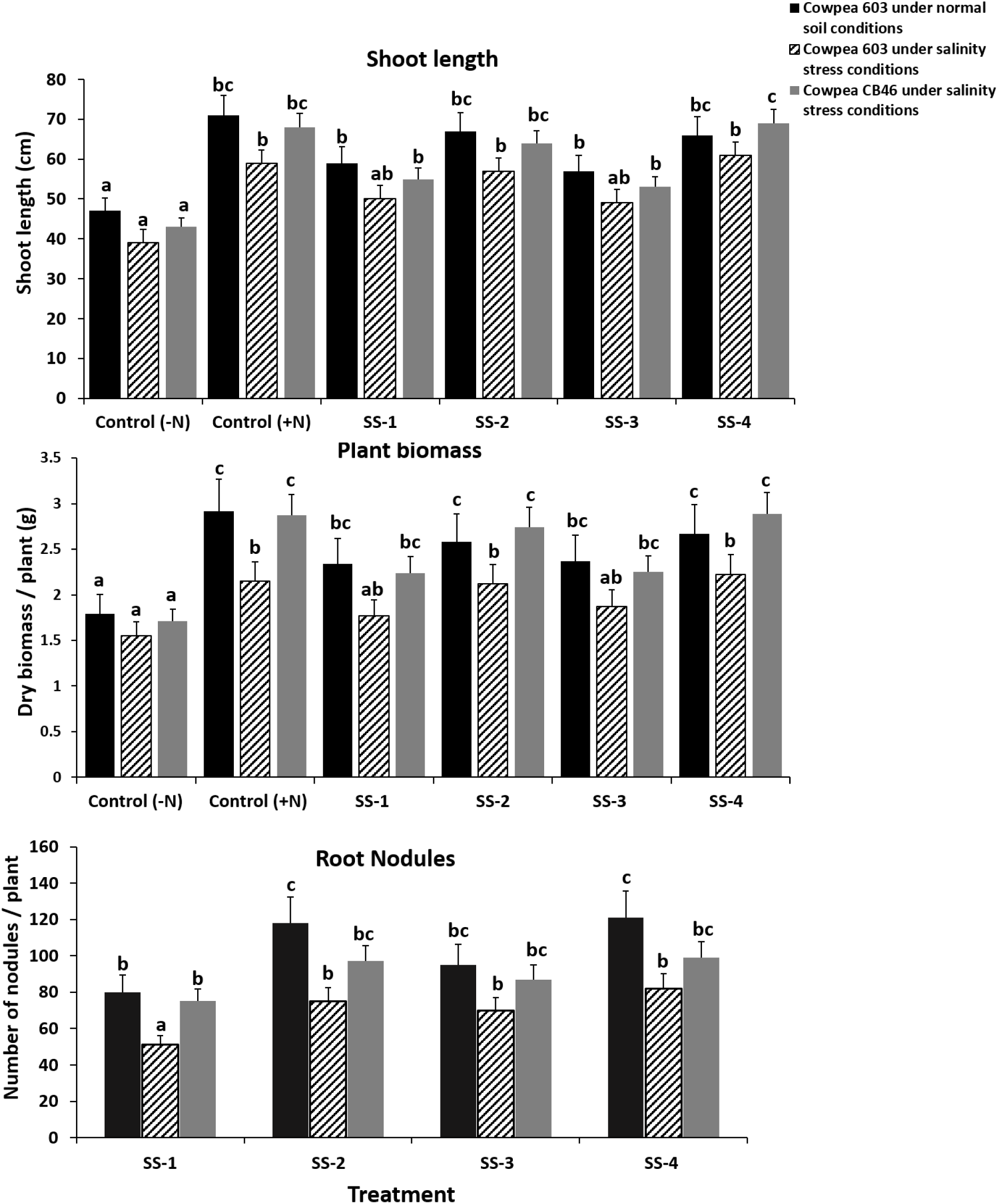
Effect of inoculation with different soil samples (SS) collected from semi-arid region of Pakistan on plant growth, (A) shoot length, (B) plant biomass and (C) number of nodules of cowpea varieties 603 and CB 46. Letters on graph bars represent statistically different values at 5% level.

The relative increase in plant biomass varied between 35-68% in treated cowpea 603 plants grown under normal soil conditions over the control (without soil inoculum). Treated cowpea 603 and CB 46 plants grown under salinity stress showed a 10-49% and 20-52% increase in plant biomass, respectively (Fig. 1; middle row). In addition, cowpea 603 and CB 46 plants inoculated with SS-2 and SS-4 soils produced more (or slightly more) nodules per plant compared to plants inoculated with SS-1 and SS-3 under normal as well as salinity-stress conditions although salinity stress did have an inhibiting effect on nodule number, which to some extent was mitigated by the CB 46 genotype (Fig. 1; bottom row).

In summary, plants inoculated with SS-2 and SS-4 showed overall better results in comparison to plants inoculated with SS-1 and SS-3 under both normal and salinity stress conditions. The differences in moisture, OM, and available mineral contents between SS-1 and SS-3 soils versus SS-2 and SS-4 soils may explain the different growth effects of the cowpea cultivars.

### eDNA analysis of SS-1 and SS-4

We chose two of the four rhizosphere soils tested above for eDNA analysis. SS-1 was an example of a soil that when added to the trap plant pot did not increase plant growth over the control; it was more saline and drier than the other soils. In contrast SS-4, the sample that resulted in growth stimulation, had the lowest electrical conductivity and the closest to neutral pH of the four soils tested.

A comparison of the microbial eDNA shows the overall pattern of bacterial sequences in the two samples (Fig. 2). Both SS-1 and SS-4 soils are dominated by Actinobacteria, Firmicutes, and Proteobacteria. The relative abundance of actinobacterial eDNA sequences was not much different between the two soil types (28.0 vs. 27.4%; SS-1 vs. SS-4) but differed slightly for the Firmicutes (28.5% vs. 26.4%) and Proteobacteria (23.6% vs. 24.9%) suggesting a possible trend. The values for Acidobacteria (6.7% vs. 7.4%) and Chloroflexi (5.5% vs. 6.8%) also varied depending on the soil sample (Table S2). In addition, eDNA relative abundances also differed between the SS-1 and SS-4 treatments. The general trend for the less-represented phyla was a reduction in DNA sequence abundance in SS-4 except for Nitrospirae, which increased. Phyla with DNA sequence abundance levels below 0.5% are also listed in Table S2.

**Fig. 2.**
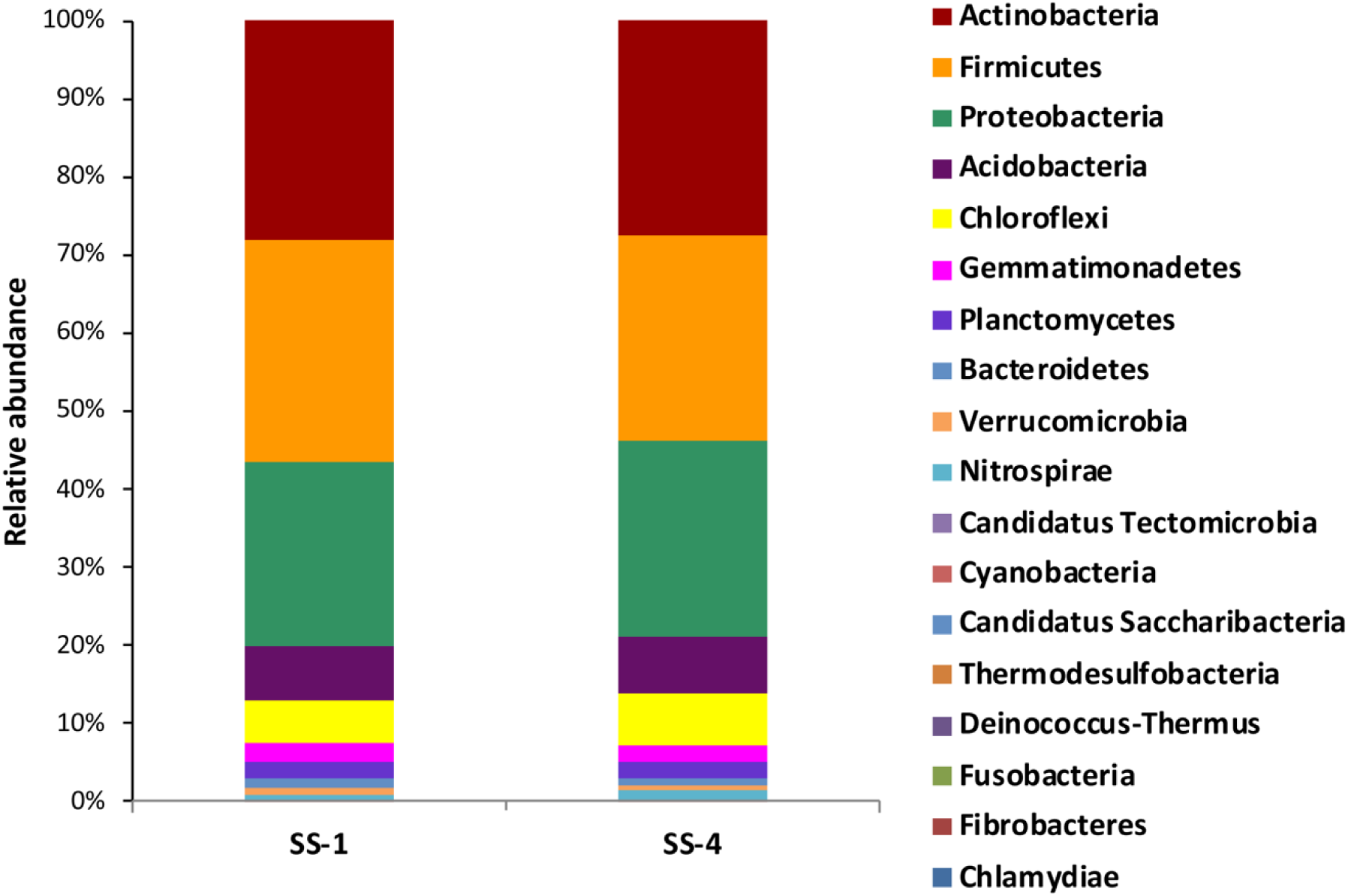
Relative abundance of eDNA sequences found in two different soils; SS-1, which was inhibitory to cowpea growth and SS-4, which promoted growth. Both soils were isolated from the Mandi Bahauddin region of Pakistan.

### Nodule microbiome analysis

Investigations of the nodule metagenomes from SS-1— SS-4 (nodule samples were designated sn1—sn4) showed that the dominant bacterial species in all the cowpea nodule microbiomes was the alpha-proteobacterial genus *Bradyrhizobium* based on relative abundance (Fig. 3; Table S3). We could not determine with certainty the identity of the strain beyond the genus level, but several different *Bradyrhizobium* species are known to nodulate cowpea in Africa (Steenkamp et al. 2004; Pule-Meulenberg et al. 2010). Detected in much lower percentages were DNA signatures of other bradyrhizobia including *Bosea*, *Afipia*, *Rhodopseudomonas*, and *Oligotropha*. Thus, the relative abundances of *Bradyrhizobiaceae* within the sn1—sn4 microbiomes were 92.3%, 84.7%, 91.5%, and 94.3%, respectively. By contrast, *Rhizobiaceae* (*Ensifer*, *Rhizobium*/*Agrobacterium*, and *Shinella*) combined with *Hyphomicrobiaceae* (*Rhodoplanes*) and *Phyllobacteriaceae* (*Mesorhizobium*) were found in the four nodule microbiomes at much lower abundances (sn1, 0.7%; sn2, 8.3%; sn3, 2.1%; and sn4, 0.3%). Other Proteobacteria detected within the nodule microbiome, albeit at very low abundance, included *Acidophilium* (Alphaproteobacteria, *Acetobacteraceae*) and *Pseudomonas* (Gammaproteobacteria, *Pseudomonadaceae*) (Table S3). The “other Proteobacteria” category exhibited a relative abundance of 3.2%, 3.8%, 3.4% and 2.8% for sn1—sn4 microbiomes. Finally, the grouping designated “non-Proteobacteria (others)” accounts for an average of 0.1% in relative abundance for sn1—sn3 and zero for sn4. These values are so low as to be undetectable in the bar graph (Fig. 3).

**Fig. 3.**
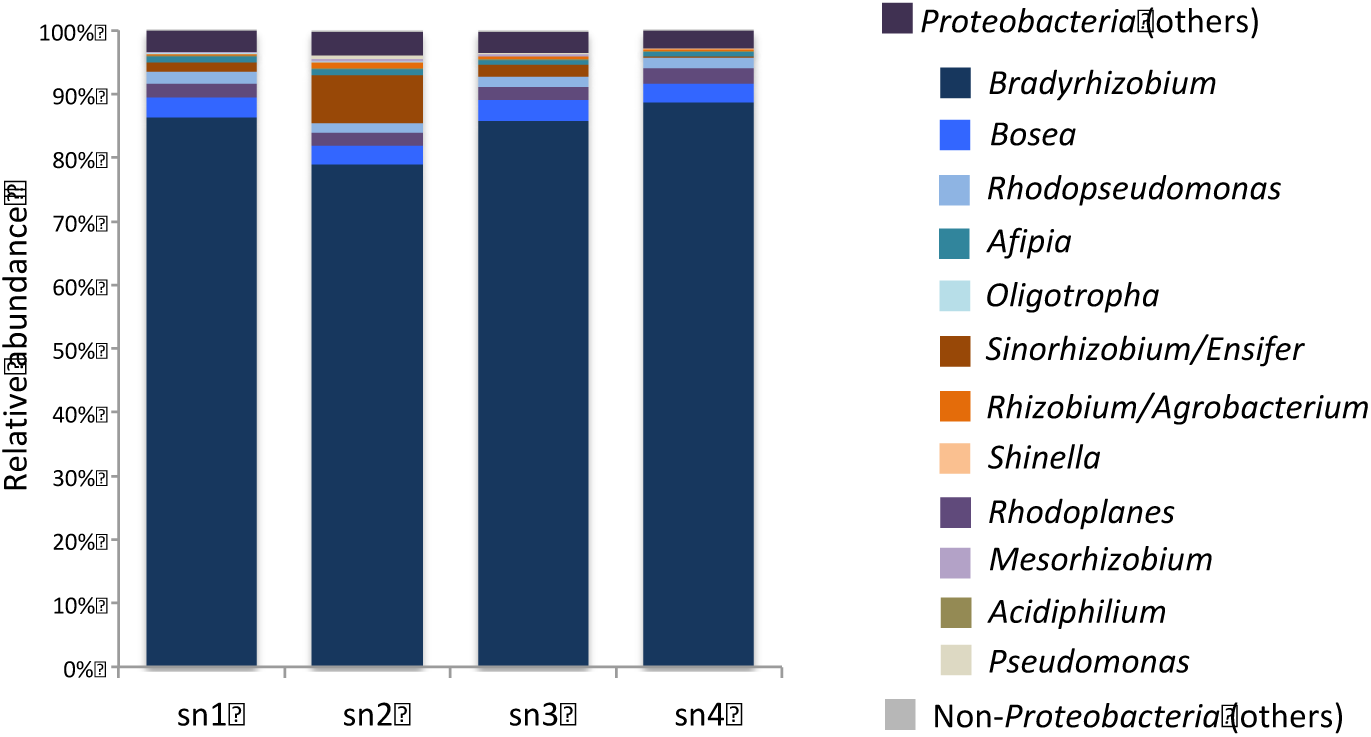
Relative abundance of microbial sequences in DNA isolated from cowpea nodules collected from four different soils from the Mandi Bahauddin region of Pakistan. In all cases, bradyrhizobia was the dominant genus, but other proteobacterial species, some of them distinct from rhizobia such as *Acidiphilium* and *Pseudomonas*, were also detected. Non-proteobacterial species, however, were barely detectable in the nodule microbiome in spite of their abundance as nodule isolates.

### Isolation and characterization of nodule bacterial isolates and their identification

A total of 51 bacterial isolates were isolated from cowpea nodules of plants grown in the four different soil samples (Table S4). Most of the isolates grew well at pH 6–8, on 1-3% NaCl, and at 28-37°C, but only 35-40% of the bacterial isolates (*Bacillus* and some *Pseudomonas* strains) grew at 4 or 42°C (data not shown).

Following a thorough morphological and physiological characterization, 34 nodule isolates were selected and identified by 16S rRNA gene analysis to the closest related species. According to the sequence match results, the bacterial strains identified from root nodules of cowpea mainly belonged to bacterial genera often shown to be plant-growth promoting rhizobacteria (PGPR) namely, *Mesorhizobium, Sinorhizobium, Bradyrhizobium*, *Paenibacillus, Bacillus, Pseudomonas* and others (Table 1). Among the PGPR were the actinomycetes *Frankia* sp DSM 45818 (CPN9), S*treptomyces galilaeus* (CPN7), and *S. griseoaurantiacus* (CPN8), which were isolated from nodules. Several isolates were identified as pathogens or nosocomial pathogens including *Klebsiella aerogenes* (CPN11 and CPN20) (formerly *Enterobacter*) (Tindal et al. 2017), *Aeronomas veronii* (CPN19) (Janda and Abbott 2010), *Pseudomonas monteilii* (CPN44) (Elomari et al. 1997), and *Enterobacter cloacae* (CPN48) (Sanders and Sanders 1997) (Table 1).

**Table 1.** Identification of bacterial isolates from cowpea nodules (CPN) based on16S rRNA gene sequence analysis.

| Isolate code | Organism identified | Accession No. | Closest type strain in NCBI data base | Sequence similarity (%) |
| --- | --- | --- | --- | --- |
| CPN1 | *Mesorhizobium* | MH491055 | *Mesorhizobium ciceri* ATCC 700744 (KX226354) | 99.45 |
| CPN2 | *Ensifer* | MH491056 | *Ensifer aridi* DSM 11282 (KR780016) | 100 |
| CPN3 | *Paenibacillus* | MH491057 | *Paenibacillus polymyxa* DSM 365 (AY359634) | 99.25 |
| CPN4 | *Paenibacillus* | MH491058 | *Paenibacillus ginsengagri* DSM 19942 (AB245383) | 99.63 |
| CPN5 | *Paenibacillus* | MH491059 | *Paenibacillus zanthoxyli* JH29 (DQ358724) | 100 |
| CPN6 | *Pseudomonas* | MH491060 | *Pseudomonas putida* ATCC 12633 (KY938109) | 100 |
| CPN7 | *Streptomyces* | MH491061 | *Streptomyces galilaeus* ATCC 14969 (KU877585) | 99.21 |
| CPN8 | *Streptomyces* | MH491062 | *Streptomyces griseoaurantiacus* NBRC 13381 (HQ850417) | 99.45 |
| CPN9 | *Frankia* | MH491063 | *Frankia* sp. DSM 45818<br>(AF063640) | 98.55 |
| CPN10 | *Bradyrhizobium* | MH491064 | *Bradyrhizobium japonicum* DES<br>122 (AF234888) | 100 |
| CPN11 | *Klebsiella* | MH491065 | *Klebsiella aerogenes* CCRC<br>10370 (KT998837) | 99.23 |
| CPN12 | *Sinorhizobium* | MH491066 | *Sinorhizobium meliloti* DSM<br>30135 (JN105982) | 99.41 |
| CPN13 | *Sinorhizobium* | MH491067 | *Sinorhizobium saheli* LMG7837<br>(NR_026096) | 99.56 |
| CPN14 | *Pseudomonas* | MH491068 | *Pseudomonas fluorescens* ATCC<br>13525 (KJ676707) | 99.74 |
| CPN15 | *Bacillus* | MH491069 | *Bacillus pumilus* NRS 272<br>(KX767153) | 99.25 |
| CPN16 | *Pseudomonas* | MH491070 | *Pseudomonas putida* DSM 291<br>(HF584918) | 99.64 |
| CPN17 | *Pseudomonas* | MH491071 | *Pseudomonas fluorescens* ATCC<br>13525 (MH191400) | 98.44 |
| CPN18 | *Pseudomonas* | MH491072 | *Pseudomonas putida* NCTC<br>10936 (KC952984) | 99.85 |
| CPN19 | *Aeromonas* | MH491073 | *Aeromonas veronii* ATCC 35624<br>(MG063196) | 100 |
| CPN20 | *Klebsiella* | MH491074 | *Klebsiella aerogenes* CCRC<br>10370 (GU459208) | 99.10 |
| CPN21 | *Bradyrhizobium* | MH603868 | *Bradyrhizobium japonicum* DSM<br>30131 (AF234888) | 99.21 |
| CPN22 | *Bacillus* | MH817487 | *Bacillus safensis* LMG 26769<br>(MH177247) | 99.07 |
| CPN27 | *Mesorhizobium* | MH817488 | *Mesorhizobium ciceri* ATCC<br>700744 (KX226354) | 99.34 |
| CPN30 | *Paenibacillus* | MH817489 | *Paenibacillus amylolyticus* NRS-290 (KR085810) | 99.28 |
| CPN35 | *Bacillus* | MH603869 | *Bacillus endophyticus* DSM<br>13796 (MH192380) | 99.54 |
| CPN36 | *Bacillus* | MH603870 | *Bacillus safensis* LMG 26769<br>(MH177247) | 99.21 |
| CPN37 | *Paenibacillus* | MH603871 | *Paenibacillus polymyxa* JCM<br>2507 (AY359634) | 99.82 |
| CPN38 | *Paenibacillus* | MH603872 | *Paenibacillus amylolyticus* NRS-290 (KR085810) | 99.34 |
| CPN41 | *Bradyrhizobium* | MH603873 | *Bradyrhizobium centrosematis*<br>LMG 29515 (NR_149804) | 100 |
| CPN43 | *Mesorhizobium* | MH603874 | *Mesorhizobium olivaresii* LMG<br>29295 (FM203301) | 99.21 |
| CPN44 | *Pseudomonas* | MH603875 | *Pseudomonas monteilii* DSM<br>11388 (KJ819574) | 99.48 |
| CPN45 | *Mesorhizobium* | MH603876 | *Mesorhizobium alhagi* HAMBI<br>3019 (KY753240) | 99.36 |
| CPN47 | *Bacillus* | MH603877 | *Bacillus megaterium* ATCC<br>14581 (HM771671) | 99.18 |
| CPN48 | *Enterobacter* | MH603878 | *Enterobacter cloacae* DSM<br>16690 (GU459208) | 99.27 |

### Plant-growth promoting abilities of bacterial strains

Isolated bacterial strains were tested for several plant-growth promoting traits including phosphate solubilization, and siderophore and HCN production. Of the 34 isolates, nineteen strains were able to solubilize phosphate with the maximum activity (142.23 μg/ml) exhibited by strain CPN35. Eight strains were positive for siderophore production, and 12 strains were positive for HCN production (data not shown).

### Antifungal activity of bacterial strains

The 34 16S rRNA gene sequence-identified bacterial isolates were tested for antifungal activity against five fungal pathogens: *Fusarium oxysporum, Fusarium solani, Curvularia* sp., *Alternaria solani*, and *Aspergillus flavus.* The details of which bacterial strains are effective against the various fungi are shown in Table S5.

### Extracellular enzymes production by bacterial isolates as mechanisms of fungal growth inhibition

The ability of bacterial strains to produce extracellular enzymes such as cellulase, lipase, chitinase, amylase, and protease was also assessed. The strains varied in their ability to produce lipase, chitinase, cellulase, amylase, and proteolytic enzymes (Table S6).

## Discussion

Because arid countries such as Pakistan, which have highly saline soils, are often the first to respond to increasing temperatures brought about by a changing climate, we pursued an investigation of the effect of salinity and aridity on the soil microbial communities that make up the cowpea rhizosphere and nodule microbiome with the goal of understanding the potential of their bacteria for being used as inoculants. Trap experiments were performed using two cowpea varieties, 603 and the more salt-tolerant CB 46. Both were inoculated with soil samples from 4 different sites in the Mandi Bahauddin area of Punjab, Pakistan. Increases in salinity are known to cause changes in soil properties such as organic matter, leaching and erosion, loss of nutrients, as well as alterations in both the quantity and composition of soil microbial communities (Canfora et al. 2012; Mukhtar et al. 2018).

Cowpea 603 and CB46 plants showed a significant increase in plant growth when inoculated with SS-2 and SS-4, even under salinity stress (10-49% and 20-52% in plant biomass, respectively) post-treatment, compared to uninoculated control plants. Previous studies have shown that the soil type had a significant influence on the microbial populations of legume nodules and ultimately plant growth (Berg and Smalla 2009; Leite et al. 2017).

Through an analysis of the nodule microbiomes of plants grown in the different soils, we found that members of the *Bradyrhizobiaceae* dominated the microbiome (78.99%-88.8% for the four microbiomes), but *Rhizobiaceae* and other Proteobacteria were also detected, albeit at significantly lower levels. However, Salter et al. (2014) observed that numerous microbes were detected in the blank controls in their experiments and thus questioned whether microbiome constituents detected in low biomass samples were contaminants rather than members of the community. For example, *Bosea* and *Afipia* (*Bradyrhizobiaceae*) were suggested to be common contaminating genera (Salter et al. 2014). Because both were observed in the cowpea nodule microbiome but were not isolated from nodules, the possibility exists that either one or both of these *Bradyrhizobiaceae* might be contaminants because their abundance values were low in all four nodule microbiome samples (*Bosea*, average: 3.3%; *Afipia*, average: 1.8%). However, many *Bradyrhizobiaceae* require long incubation times before colonies become apparent on plates, and in most cases, the cowpea nodule isolate incubations in our study did not exceed one week. For many of the slow-growing *Bradyrhizobiaceae* (*Bosea*. *Rhodopseudonomas*, *Afipia*, and others), longer time periods (at least 2 weeks) are often needed for growth after isolation from nodules and transfer to culture medium. However, a phylogenetic analysis (Fig. 4) of the bacterial root nodule isolates of cowpea 603 and CB46 plants inoculated with the saline soil samples demonstrated that many were members of either *Rhizobiaceae* or *Bradyrhizobiaceae* strongly suggesting that these genera are not contaminants.

**Fig. 4.**
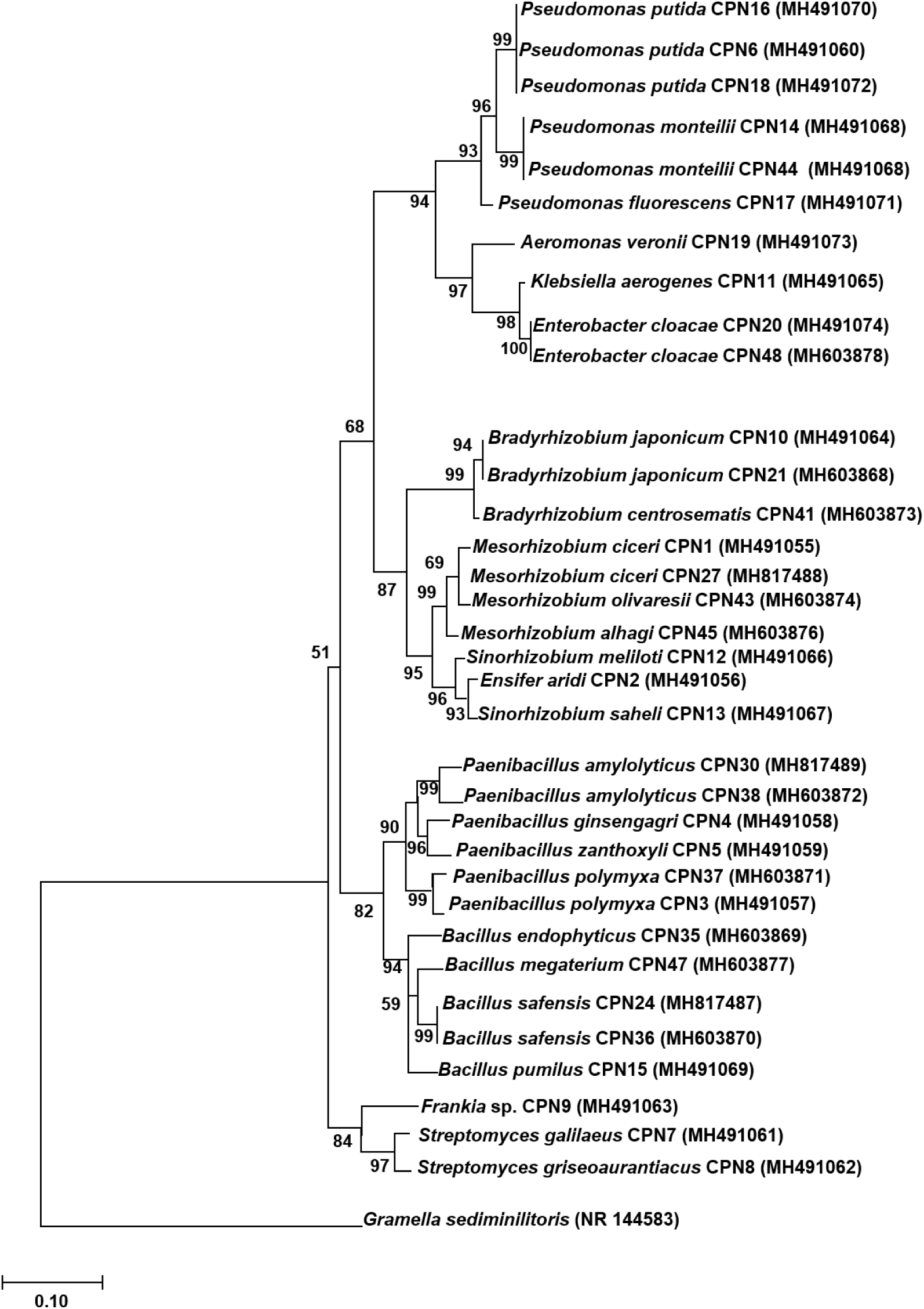
Neighbor-joining tree based on 16S rRNA gene sequences of bacterial strains isolated from cowpea nodules. The percentage of replicate trees in which the associated taxa clustered together in the bootstrap test (1,000 replicates) is shown next to the branches. The evolutionary distances were computed using the Maximum Composite Likelihood method and are in the units of number of base substitutions per site.

In any case, we do not know the reasons for the difference in relative abundance between the rhizobial data obtained from the microbiome analyses and the isolation methodology. Studies have shown that soil physicochemical properties or physiological changes within the nodule itself may affect the survivability or distribution of indigenous rhizobia isolated from cowpea nodules (Pule-Meulenberg et al. 2010; Damaris et al. 2017; Leite et al. 2017). Also, media composition, specific plant growth conditions, as well as incubation time and temperature all play a role as well as variations in the methods of microbiome analysis.

With regard to the firmicutes, the bacterial isolates investigated in this study were mostly identified as *Paenibacillus* and *Bacillus* based on isolation of bacteria from nodules (Table 1, Fig. 4). Nodule isolates from cowpea plants grown in semi-arid areas of Brazil were also known to be high in Firmicutes (*Bacillus* and *Paenibacillus*) (Costa et al. 2013), and a number of studies have reported that the majority of isolates from cowpea nodules belong to the Firmicutes, Proteobacteria, and Actinobacteria (Jaramillo et al. 2013; Leite et al. 2017). However, the metagenomic analysis of the nodule microbiome in our study demonstrated that DNA from Firmicutes and Actinobacteria was not detected in cowpea nodules; only the DNA of Proteobacteria This result may be related to the type of DNA isolation kit used or sample source/treatment (Mackenzie et al., 2015) or as yet unknown reasons. Nevertheless, both Firmicutes and Actinobacteria were isolated from surface-sterilized nodules, and our results and those of others demonstrate that these two groups are well-represented among the nodule isolates. For example, both *Streptomyces* and *Frankia* (Table 1) were isolated from cowpea nodules in this study and in other nodule studies (Tani et al. 2003; Leite et al. 2017) but were not detected in the nodule microbiome analysis.

An exceptionally low relative abundance level of *Pseudomonas* sequences was also detected in the nodule microbiome, but Gammaproteobacteria (*Pseudomonas*) and (*Enterobacter*/*Klebsiella*) were consistently isolated and characterized from cowpea nodules collected from plants grown under salt stress conditions (Table 1, Fig. 4).

In reference to the rhizosphere, an analysis of the SS-1 (growth-inhibiting) and SS-4 (growth-promoting) rhizosphere soils demonstrated the former was more saline and drier than the latter. These features as well as the reduced levels of available P, K, Ca, and Mg as well as organic matter and higher sodicity values of the SS-1 compared to SS-4 soil most likely explain the lack of fertility in SS-1 soil as well as its reduced effect on plant growth.

## Summary

Numerous bacteria were detected and isolated from the SS-1 and SS-4-grown cowpea nodules. Some were found to promote plant growth directly by nitrogen fixation and mineral solubilization, while others indirectly by production of antibacterial and antifungal compounds (Malik et al. 1997; Gupta et al. 2015; Goswami et al. 2016). Many of the isolates obtained in this study were positive for more than two PGPR traits. Several of the strains were likely to have nitrogen fixation ability and greater than 55% of bacterial isolates from cowpea nodules exhibited phosphate-solubilizing ability. Finally, the majority of the *Pseudomonas, Bacillus,* and *Enterobacter* strains isolated from cowpea nodules were positive for siderophore and HCN production. Our study also showed that isolates belonging to *Pseudomonas*, *Paenibacillus,* and *Bacillus* produced antifungal compounds against different fungal pathogens, including *Fusarium oxysporum, Fusarium solani, Curvularia* sp. Both *Pseudomonas* and *Bacillus* have been reported as growth inhibitors for different fungal pathogens and provide protection against a number of plant diseases (Ali et al. 2015; Shahid et al. 2017; Khan et al. 2017). More than one-half of the isolated strains especially members of *Paenibacillus, Bacillus, Pseudomonas* and *Enterobacter*, exhibited cellulase, chitinase, amylase, and protease activity, and a large number of PGPR strains (*Pseudomonas*, *Enterobacter, Rhizobium* and *Bacillus*) promote plant growth and suppress plant diseases by producing siderophores, hydrolytic enzymes, and HCN (Mehnaz et al. 2010; Chen et al. 2013; Khan et al. 2018).

This study aimed to assess the effect of native soil microbial communities on the composition of the nodule microbiome in cowpea under salinity stress conditions. The results of trap experiments confirmed that cowpea inoculated with two of the soil samples from Mandi Bahauddin in Punjab, Pakistan displayed enhanced growth as compared to the uninoculated negative control group. Insight into the bacterial communities of rhizosphere soil and cowpea root nodules, which served as trap plants was obtained. Although the eDNA of nodule microbiome consisted of mainly of Proteobacteria, various other bacteria including *Paenibacillus*, *Bacillus* and several Actinobacteria, which were isolated from cowpea nodules inoculated with one of the four different salinity-stressed soils. The bacterial strains isolated from root nodules have potential for stimulating plant growth and development and several have been characterized as PGPR. Taken together, these bacterial strains may serve as useful biofertilizers and plant growth promoters for crops growing in salinity-affected areas. Whether these bacteria will restore soil fertility in the dry arid soils of certain regions of Mandi Bahauddin will be a focus of future research.

## Supporting information

All Supplementary Files

## Acknowledgments

We are grateful to the Higher Education Commission, Pakistan for an IRSIP fellowship at the Department of Molecular, Cell and Developmental Biology, University of California Los Angeles, U.S.A. A Community Sequence Project (CSP 1571) from the U.S. Department of Energy Joint Genome Institute (DOE/JGI) also funded the project. Work conducted by the U.S. Department of Energy Joint Genome Institute, a DOE Office of Science User Facility, is supported by the Office of Science of the U.S. Department of Energy under Contract No. DE-AC02-05CH11231. Additional funding for studying arid soils came from the Shanbrom Family Foundation to AMH. We also acknowledge the contributions of Samantha Lieberman and other members of the Hirsch lab (UCLA) for their help and support with various experiments and are grateful to Prof. T. Close of the University of California, Riverside for seeds of *V. unguiculata* CB 46.

## Author contributions

All authors contributed to the development, writing, and completion of this manuscript. A first draft of the manuscript was submitted as part of the Ph.D. degree research of Salma Mukhtar at Forman Christian College (A Charter University) in Lahore, Pakistan, under the direction of Prof. Kauser Malik and Dr. Samina Mehnaz.

## References

1. Adviento-Borbe, M.A., Doran, J.W., Drijber, R.A., Dobermann, A. 2006. Soil electrical conductivity and water content affect nitrous oxide and carbon dioxide emissions in intensively managed soils. J. Environ. Qual. 35: 1999–2010.

2. Ali, G.S., Norman, D., El-Sayed, A.S. 2015. Soluble and volatile metabolites of plant growth-promoting rhizobacteria (PGPRs): role and practical applications in inhibiting pathogens and activating induced systemic resistance (ISR). Adv. Bot. Res. Acad. Press 75: 241–284.

3. Ali, Y., Aslam, Z., F. Hussain, Shakur, A. 2004. Genotype and environmental interaction in cowpea (*Vigna unguiculata* L.) for yield and disease resistance. Int. J. Environ. Sci. Technol. 1: 119–123

4. Anderson, J.M., Ingram, J.S. 1993. Tropical Soil Biology and Fertility: A Handbook of Methods. pp. 93–94. 2nd ed. CAB International, Wallingford, UK.

5. Ardley, J.K., Parker, M.A., O’Hara, G.W., Reeve, W.G., Yates R.J., Dilworth, M.J., Willems, A., Howieson, J.G. 2012. *Microvirga lupini* sp. nov., *Microvirga lotononidis* sp. nov. and *Microvirga zambiensis* sp. nov. are alphaproteobacterial root-nodule bacteria that specifically nodulate and fix nitrogen with geographically and taxonomically separate legume hosts. Int. J. Syst. Evol Microbiol. Nov;62(Pt 11):2579–88. doi: 10.1099/ijs.0.035097-0.

6. Aserse, A.A., Räsänen, L.A., Aseffa, F., Hailemariam, A., Lindström, K. 2013. Diversity of sporadic symbionts and nonsymbiotic endophytic bacteria isolated from nodules of woody, shrub, and food legumes in Ethiopia. Appl. Microbiol. Biotechnol. 97: 10117–10134.

7. Beijerinck, M.W. 1888. The root-nodule bacteria. Bot. Zeitung. 46: 725–804.

8. Berg, G., and Smalla, K. 2009. Plant species and soil type cooperatively shape the structure and function of microbial communities in the rhizosphere. FEMS Microbiol. Ecol. 68: 1–13.

9. Bontemps, C., Elliott, G.N., Simon, M.F., Dos Reis, F.B., Gross, E., Lawton, R.C., Neto, N.E., de Fátima Loureiro, M., De Faria, S.M., Sprent, J.I., James, E.K., Young, J.P. 2010. *Burkholderia* species are ancient symbionts of legumes. Mol. Ecol. 19:44–52..

10. Canfora, L., Bacci, G., Pinzari, F., Lo Papa, G., Dazzi, C., Benedetti, A. 2014. Salinity and bacterial diversity: to what extent does the concentration of salt affect the bacterial community in a saline soil? PLoS One. 9:e106662.

11. Chen, L.H., Lin, C.H., Chung, K.R. 2013. A nonribosomal peptide synthetase mediates siderophore production and virulence in the citrus fungal pathogen *Alternaria alternata*. Mol. Plant Path. 14: 497–505.

12. Costa, E.M., Nóbrega, R.S.A., Carvalho, F., Trochmann, A., Ferreira, L.V.M., Moreira, F.M.S. 2013. Plant growth promotion and genetic diversity of bacteria isolated from cowpea nodules. Pesq. Agropec. Bras. 48: 1275–1284.

13. Damaris, K.O., Nyaboga, E.N., Wagacha J.M., Mwaura, F.B. 2017. Morphological and genetic diversity of Rhizobia nodulating cowpea (*Vigna unguiculata* L.) from agricultural soils of lower eastern Kenya. Int. J. Microbiol. 2017: 8684921.

14. Deng, Z.S., Zhao, L.F., Kang, Z.Y., Yang, W.Q., Lindström, K., Wang, I.T., Wei, G.H. 2011. Diversity of endophytic bacteria within nodules of the *Sphaerophysa salsula* in different regions of Loess Plateau in China. FEMS Microbiol. Ecol. 76: 463–475.

15. DeSantis, T.Z., Hugenholtz, P., Larsen, N., Rojas, M., Brodie, E.L., Keller, K., Huber, T., Dalevi, D., Hu, P., Andersen, G.L. 2006. Greengenes, a chimera-checked 16S rRNA gene database and workbench compatible with ARB. Appl. Environ. Microbiol. 72:5069–72.

16. Eberl, L., Vandamme. P. 2016. Members of the genus *Burkholderia*: good and bad guys [version 1; peer review: 3 approved]. F1000Research 2016, 5(F1000 Faculty Rev):1007 (https://doi.org/10.12688/f1000research.8221.1)

17. Elomari, M. Coroler, L., Verhille, S., Izard, D., Leclerc, H. 1997. *Pseudomonas monteilii* sp. nov., isolated from clinical specimens. Int. J. Syst. Bacteriol. 47 (3): 846–852. doi:10.1099/00207713-47-3-846.

18. Estrada-de los Santos, P., Palmer, M., Chávez-Ramírez, B., Beukes, C., Steenkamp, E.T., Briscoe, L., Khan, N., Maluk, M., Lafos, M., Humm, E., Arabit, M., Crook, M., Gross, E., Simon, M.F., Bueno dos Reis Junior, F., Whitman, W.B., Shapiro, N., Poole, P.S., Hirsch, A.M., Venter, S.N., James, E.K. 2018. Whole genome analyses suggest that *Burkholderia sensu lato* contains two further novel genera in the “rhizoxinica-symbiotica group” (*Mycetohabitans* gen. nov., and *Trinickia* gen. nov.): implications for the evolution of diazotrophy and nodulation in the *Burkholderiaceae*. Genes. 9(8), 389; https://doi.org/10.3390/genes9080389.

19. Garg, N, Geetanjali. 2007. Symbiotic nitrogen fixation in legume nodules: process and signaling. A review. Agron. Sustain. Dev. 27: 59–68.

20. Goswami, D., Thakker, J.N., Dhandhukia P.C. 2016. Portraying mechanics of plant growth promoting rhizobacteria (PGPR): A review. Cog. Food Agri. 2: 11275.

21. Graham, P.H. 2008. Ecology of the root-nodule bacteria of legumes, in Nitrogen-fixing Leguminous Symbioses, eds. M.J. Dilworth, E.K. James, J. Sprent I, and W.E. Newton (Dordrecht: Springer), pp: 23–58.

22. Gupta, G., Parihar, S.S., Ahirwar, N.K., Snehi, S.K., Singh, V. 2015. Plant growth promoting rhizobacteria (PGPR): current and future prospects for development of sustainable agriculture. J. Microb. Biochem. Technol. 7: 96–102.

23. Gyaneshwar, P., Hirsch, A.M., Moulin, L., Chen, W.M., Elliott, G.N., Bontemps, C., Estrada-de los Santos, P., Gross, E., Dos Reis, F.B., Sprent, J.I., Young, J.P.W., James, E.K. 2011. Legume-nodulating betaproteobacteria: Diversity, host range, and future prospects. Mol. Plant-Microbe Interact. 24: 1276–1288.

24. Helms, D., Panella, L., Buddenhagen, L.W., Tucker, C.L., Gepts, P.L. 1991. Registration of ‘California Blackeye 46’ cowpea. Crop Sci. 331:1703.

25. Hirsch, A.M. 1992. Developmental biology of legume nodulation. New Phytologist 122: 211–237.

26. Imran, M., Qamar, I.A., Muhammad, S., Mahmood, I.A., Chathha, M.R., Gumani, Z.A., Anjum, A.S., Shahid, M.N. 2012. Comparison of different cowpea varieties/lines for green fodder and grain yield under rainfed conditions of Islamabad, Pakistan. Sarhad J. Agric. 28: 41–46.

27. Janda, J.M., Abbott, S.L. 2010. The genus *Aeromonas*: Taxonomy, pathogenicity, and infection. Clin. Microbiol. Rev. 23:35–75.

28. Jaramillo, P.M.D., Guimarães, A.A., Florentino, L.A., Silva, K.B., Nóbrega, R.S.A., Moreira, F.M.S. 2013. Symbiotic nitrogen-fixing bacterial populations trapped rom soils under agroforestry systems in the Western Amazon. Sci. Agric. 70: 397–404.

29. Kasana, R.C., Salwan, R., Dhar, H., Dutt, S., Gulati, A. 2008. A rapid and easy method for the detection of microbial cellulases on agar plates using gram’s iodine. Curr Microbiol. 57(5): 503–507.

30. Kaur, K., Dattajirao, V., Shrivastava, V., Bhardwaj, U. 2012. Isolation and characterization of chitosan-producing bacterial from beaches of Chennai, India. Enzy. Res. 2012: 421683.

31. Khan, N., Martínez-Hidalgo, P., Ice, T.A., Maymon, M., Humm, E.A., Nejat, N., Sanders, E.R., Kaplan, D., Hirsch, A.M. 2018. Antifungal activity of *Bacillus* species against *Fusarium* and analysis of the potential mechanisms used in biocontrol. Front. Microbiol. 9: 2363.

32. Khan, N., Maymon, M., Hirsch, A.M. 2017. Combating *Fusarium* infection using *Bacillus*-based antimicrobials. Microorganisms 5: E75.

33. Kuddus, M., Ahmad, I.Z. 2013. Isolation of novel chitinolytic bacteria and production optimization of extracellular chitinase. J. Gen. Eng. Biotech. 11: 39–46.

34. Kumar, S., Stecher, G., Tamura, K. 2016. MEGA7: Molecular Evolutionary Genetics Analysis version 7.0 for bigger datasets. Mol Biol Evol. 33: 1870–1874.

35. Kumar, K.V., Srivastava, S., Singh, N., Behl, H.M. 2009. Role of metal resistant plant growth promoting bacteria in ameliorating fly ash to the growth of *Brassica juncea*. J. Haz. Mat. 170:51–57.

36. Lafay, B., Burdon, J.J. 2007. Molecular diversity of legume root-nodule bacteria in Kakadu National Park, Northern Territory, Australia. PLoS One 2(3): e277.

37. Langille, M.G., Zaneveld, J., Caporaso, J.G., McDonald, D., Knights, D., Reyes, J.A, et al. 2013. Predictive functional profiling of microbial communities using 16S rRNA marker gene sequences. Nat. Biotechnol. 31:814–21.

38. Langmead B, Trapnell C, Pop M, Salzberg SL. 2009. Ultrafast and memory-efficient alignment of short DNA sequences to the human genome. Genome Biol 10: R25.

39. Leite, J., Fischer, D., Rouws, L.F.M., Fernandes-Júnior, P.I., Hofmann, A., Kublik, S., Schloter, M., Xavier, G.R., Radl, V. 2017. Cowpea nodules harbor non-rhizobial bacterial communities that are shaped by soil type rather than plant genotype. Front. Plant Sci. 7: 2064.

40. Mackenzie, B.W., Waite, D.W., Taylor, M.W. 2015. Evaluation variation in human gut microbiota profiles due to DNA extraction method and inter-subject differences. Front. Microbiol. 6:130.

41. Malik, K.A., Bilal, R., Mehnaz, S., Rasool, G., Mirza, M.S. Ali, S., 1997. Association of nitrogen-fixing, plant growth promoting rhizobacteria (PGPR) with kallar grass and rice. Plant Soil. 194: 37–44

42. Martínez-Hidalgo, P and Hirsch, A.M. 2017. The nodule microbiome: N_2_-fixing rhizobia do not live alone. Phytobiomes J. 1: 70–82.

43. Mehnaz, S., Baig, D.N., Lazarovits, G. 2010. Genetic and phenotypic diversity of plant growth promoting rhizobacteria isolated from sugarcane plants growing in Pakistan. J. Microbiol. Biotechnol. 20: 1614–1623.

44. Muhammad, D., A. Hussain, S. Khan and M.B. Bhatti. 1993. Variability for green fodder yield and quality in cowpea under rainfed conditions. Pak. Agric. Res. 14: 154–158.

45. Mukhtar, S., Mirza, B.S., Mehnaz, S., Mirza, M.S., Mclean, J., Malik, K.A. 2018. Impact of soil salinity on the microbial structure of halophyte rhizosphere microbiome. World J Microbiol Biotechnol. 34(9): 136.

46. Olsen, S.R., Cole, C.V., Watanabe, F.S., Dean, L.A. 1954. Estimation of available phosphorus in soils by extraction with sodium bicarbonate. USDA Circular 939:1–19. Gov. Printing Office Washington D.C.

47. Perez-Miranda, S., Cabirol, N., George-Tellez, R., Zamudio-Rivera, L.S., Fernandez, F.J. (2007). O-CAS, a fast and universal method for siderophore detection. J. Microbiol. Methods 70: 127–131.

48. Pikovskaya, R. 1948. Mobilization of phosphorus in soil in connection with vital activity of some microbial species. Mikrobiologiya 17: 362–370.

49. Pruitt, K.D., Tatusova, T., Maglott, D.R. 2005. NCBI Reference Sequence (RefSeq): a curated non-redundant sequence database of genomes, transcripts and proteins. Nucleic Acids Res. 33(Database issue):D501–4.

50. Pule-Meulenberg, F., Belange, A.K., Krasova-Wade, T., Dakora, F.D. 2010. Symbiotic functional and bradyrhizobial biodiversity of cowpea (*Vigna unguiculata* L. Walp.) in Africa. BMC Microbiol. 10:89.

51. Sadasivam, S., Manickam, A. 1992. Biochemical Methods for Agricultural Sciences. Wiley Eastern Ltd, New Delhi, p. 246.

52. Saitou N, Nei M. 1987. The neighbor-joining method: A new method for reconstructing phylogenetic trees. Mol. Biol. Evol. 4: 406–425.

53. Sanders, W.E. Jr., Sanders. C.C. 1997. *Enterobacter* spp.: pathogens poised to flourish at the turn of the century. Clin. Microbiol. Rev.10,220–241.

54. Salter, S.J., Cox, J.M., Turek, E.M., Calus, S.T., Cookson, W.O., Moffatt, M.F., Turner, P., Parkhill, J., Lornan, N.J., Walker, A.W. 2014. Reagent and laboratory contamination can critically impact sequenced-based microbiome analyses. BMC Biol. 12:87.

55. Sawada, H., Kuykendall, L.D., Young, J.M. 2003. Changing concepts in the systematics of bacterial nitrogen-fixing legume symbionts. J. Gen. Appl. Microbiol. 49: 155–179.

56. Sawana A, Adeolu M, Gupta RS. 2014. Molecular signatures and phylogenomic analysis of the genus *Burkholderia*: Proposal for division of this genus into the emended genus *Burkholderia* containing pathogenic organisms and a new genus *Paraburkholderia*, gen. nov. harboring environmental species. Front. Genet. 5: 429. doi:10.3389/fgene.2014.00429. PMC 4271702.

57. Schwartz, A.R., Ortiz, I., Maymon, M., Herbold, C.W., Fujishige, N.A, Vijanderan, J.A., Villella, W., Hanamoto, K., Diener, A., Sanders, E.R., DeMason, D.A., Hirsch, A.M. 2013. *Bacillus simplex*—a little known PGPB with anti-fungal activity—alters pea legume root architecture and nodule morphology when coinoculated with *Rhizobium leguminosarum* bv. *viciae*. Agronomy. 3:595–620; doi:10.3390/agronomy3040595.5.

58. Shahid, I., Rizwan, M., Baig, D.N., Saleem, R.S., Malik, K.A., Mehnaz, S. 2017. Secondary metabolites production and plant growth promotion by *Pseudomonas chlororaphis* subsp. *aurantiaca* strains isolated from cotton, cactus and para grass. J Microbiol Biotechnol 27: 480–491.

59. Shi, B., Chang, M., Martin, J., Mitreva, M., Lux, R., Klokkevold, P., Sodergren, E., Weinstock, G.M., Haake, S.K., Li, H. 2015. Dynamic changes in the subgingival microbiome and their potential for diagnosis and prognosis of periodontitis. mBio 6(1):e01926–14.

60. Sierra, G. 1957. A simple method for the detection of lipolytic activity of micro-organisms and some observations on the influence of the contact between cells and fatty acid substrates. A. Van. Leeuw. J. Microbiol. 23: 15–22.

61. Sigmon, J. 2008. The starch hydrolysis test. http://www.asmscience.org/content/education/imagegallery/image.3172.

62. Steenkamp, E.T., Stepkowski, T., Przymusiak, A., Bothaa, W.J., Law, I.J. 2008. Cowpea and peanut in southern Africa are nodulated by diverse *Bradyrhizobium* strains harboring nodulation genes that belong to the large pantropical clade common in Africa. Mol. Phylo.. Evol. 48:1131–1144.

63. Sturz, A., Christie, B., Matheson, B., and Nowak, J. 1997. Biodiversity of endophytic bacteria which colonize red clover nodules, roots, stems and foliage and their influence on host growth. Biol. Fert. Soils 25: 13–19.

64. Sulieman, S., Tran, L.P. 2014. Symbiotic nitrogen fixation in legume nodules: metabolism and regulatory mechanisms. Int J Mol Sci. 15(11): 19389–19393.

65. Tamura, K., Nei, M., Kumar, S., 2004. Prospects for inferring very large phylogenies by using the neighbor-joining method. Proc. Natl. Acad. Sci. USA. 101: 11030–11035.

66. Tani, C, Sasakawa, H, Takenouchi, K. 2003. Isolation of endophytic *Frankia* from root nodules of *Casuarina equisetifolia* and infectivity of the isolate to host plants. Soil Sci. Plant Nutr, 49: 137–142.

67. Tindal, B.J., Sutton, G., Garrity, G.M. 2017, *Enterobacter aerogenes* Hormaeche and Edwards 1960 (Approved Lists 1980) and *Klebsiella mobilis* Bascomb et al. 1971 (Approved Lists 1980) share the same nomenclatural type (ATCC 13048) on the approved lists and are homotypic synonyms, with consequences for the name *Klebsiella mobilis* Bascomb et al. 1971 (Approved Lists 1980). Int. J. Syst. Evol. Microbiol. 67:502–504.

68. Velázquez, E., Martínez-Hidalgo, P., Carro, L., Alonso, P., Peix, A., Trujillo, M. E., and Martínez-Molina, E. 2013. Nodular endophytes: An untapped diversity, Beneficial Plant Microbial Interactions: Ecology and Applications. CRC Press, Boca Raton, FL. pp: 215–236.

69. Walkley, A., Black, I.A. 1934. An examination of Degtjareff method for determining soil organic matter and a proposed modification of the chromic acid titration method. Soil Sci. 37:29–37.

70. Watanabe, F., Olsen, S., 1965. Test of an ascorbic acid method for determining phosphorus in water and NaHCO3extracts from soil 1. Soil Sci. 29: 677–678.

71. Weisburg, W.G., Barns, S.M., Pelletier, D.A., Lane, D.J. 1991. 16S ribosomal DNA amplification for phylogenetic study. J. Bact. 173: 697–703.

72. Xu, L., Zhang, Y., Wang, L., Chen, W., and Wei, G. 2014. Diversity of endophytic bacteria associated with nodules of two indigenous legumes at different altitudes of the Qilian Mountains in China. Syst. Appl. Microbiol. 37: 457–465.

