## Supplementary material for "Impact of soil salinity on the cowpea nodule-microbiome and the isolation of halotolerant PGPR strains to promote plant growth under salinity stress": All Supplementary Files

Supplementary Figures

Fig. S1. Geological information of sample area in Mandi Bahauddin and locations of the four regions where the soil sample were taken.

Fig S2. Comparison of plant growth between inoculated and uninoculated control plants; (A) Cowpea 603 under normal soil conditions, (B) Cowpea 603 under salinity-stress conditions and (C) Cowpea CB46 under salinity-stress conditions

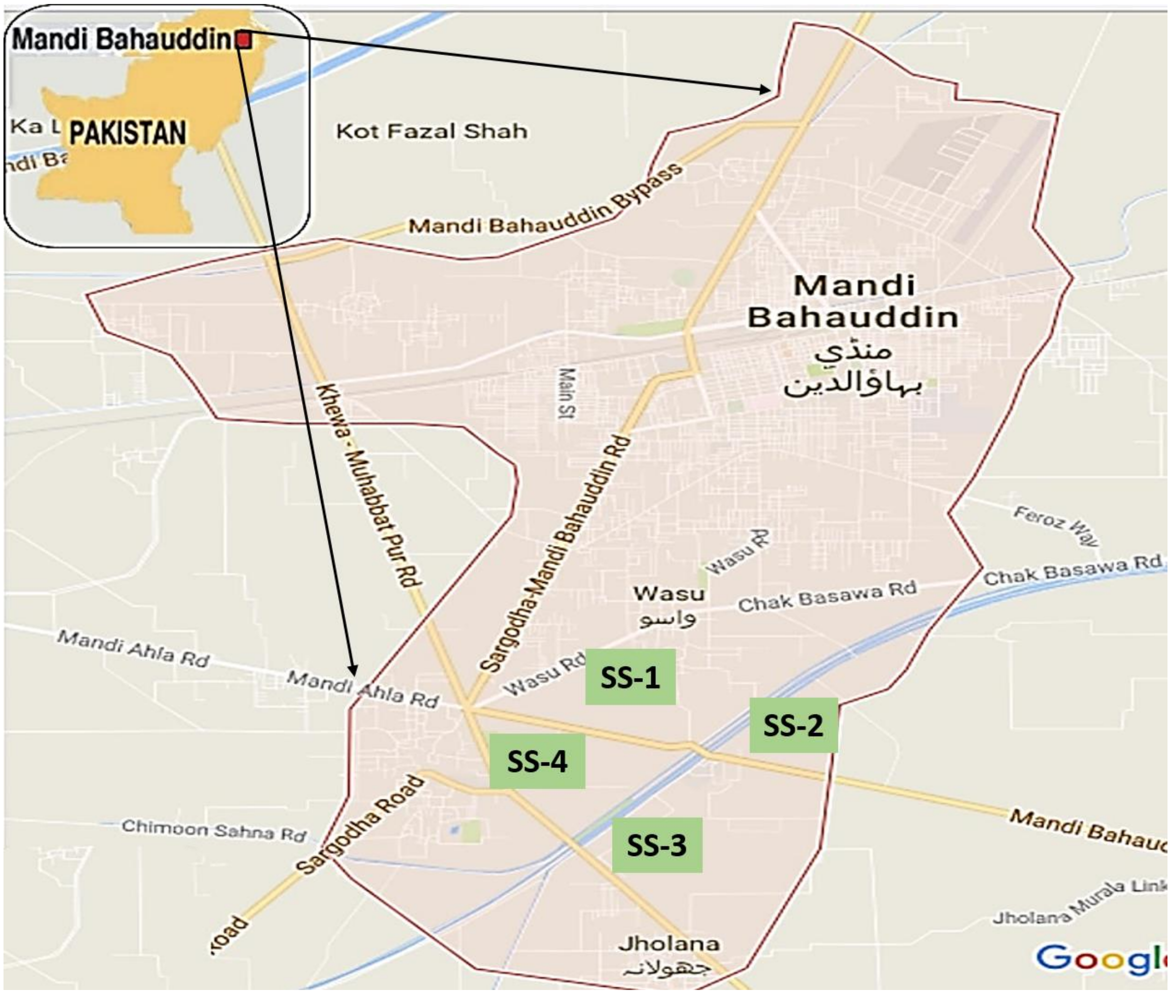


**Figure S1.** Geological information of sampling area, Mandi Bahauddin, Pakistan

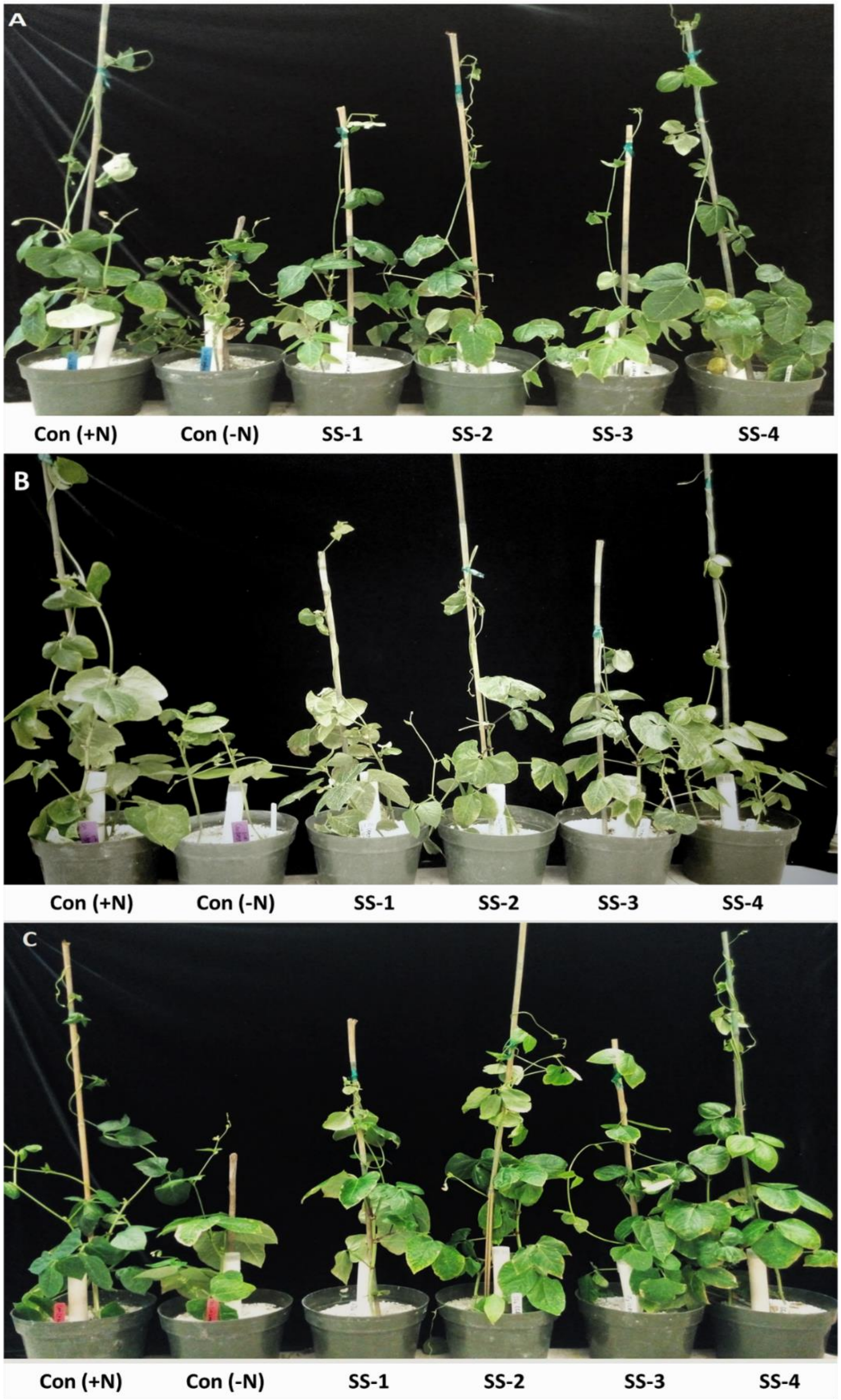
A
Con (+N)
Con (-N)
SS-1
SS-2
SS-3
SS-4
B
Con (+N)
Con (-N)
SS-1
SS-2
SS-3
SS-4
C
Con (+N)
Con (-N)
SS-1
SS-2
SS-3
SS-4

**Figure S2.**

**Table S1. Physicochemical characteristics of cowpea rhizospheric soil samples collected from different locations of semiarid regions in Pakistan.**

| Soil physicochemical characteristics | Soil sample 1 (SS-1) | Soil sample 2 (SS-2) | Soil sample 3 (SS-3) | Soil sample 4 (SS-4) |
|---|---|---|---|---|
| $EC_{1:1}$ (dS/m) | $5.05^{ab}$ | $4.85^{ab}$ | $5.29^{b}$ | $4.56^{a}$ |
| pH | $7.89^{ab}$ | $7.65^{a}$ | $8.11^{b}$ | $7.59^{a}$ |
| Temperature (°C) | $39.24^{b}$ | $38.31^{a}$ | $38.89^{ab}$ | $38.54^{a}$ |
| Moisture (%) | $18^{a}$ | $19^{a}$ | $24^{b}$ | $21^{ab}$ |
| Texture class | Silty loam | Sandy loam | Sandy loam | Sandy loam |
| OM ($g.Kg^{-1}$) | $24.56^{a}$ | $31.75^{b}$ | $21.23^{a}$ | $29.68^{ab}$ |
| P ($mg.kg^{-1}$) | $12.91^{a}$ | $15.61^{ab}$ | $21.37^{b}$ | $11.01^{a}$ |
| K ($mg.kg^{-1}$) | $0.51^{b}$ | $0.36^{a}$ | $0.45^{ab}$ | $0.39^{a}$ |
| Ca ($mg.kg^{-1}$) | $1.39^{ab}$ | $1.46^{b}$ | $1.38^{ab}$ | $1.27^{a}$ |
| Mg ($mg.kg^{-1}$) | $1.05^{a}$ | $1.21^{b}$ | $0.98^{a}$ | $1.15^{ab}$ |
| $NO^{-3}$ ($mg.kg^{-1}$) | $12.36^{ab}$ | $11.12^{a}$ | $10.69^{a}$ | $12.85^{ab}$ |
| H+Al ($mg.kg^{-1}$) | $62.27^{b}$ | $53.54^{a}$ | $56.24^{a}$ | $59.69^{ab}$ |
| V ($mg.kg^{-1}$) | $4.03^{a}$ | $3.57^{a}$ | $4.26^{b}$ | $4.16^{ab}$ |
| CEC ($mg.dm^{-3}$) | $61.45^{ab}$ | $67.52^{b}$ | $58.98^{a}$ | $64.36^{ab}$ |
| SAR | $12.42^{b}$ | $10.41^{a}$ | $11.18^{ab}$ | $11.47^{ab}$ |

Note: EC (Electrical conductivity); OM (Organic matter); P (Phosphorous); K (Potassium); Ca (Calcium); Mg (Magnesium); $NO^{-3}$ (Nitrate ion); H+Al (potential acidity); V (base saturation index); CEC (Cation exchange capacity) and SAR (*Sodium adsorption ratio*)

*Values are mean of three samples (mean ± SE)

**Table S2.** Percent relative abundance of phyla in SS-1 and SS-4

| Phylum | SS-1 | SS-4 |
|---|---|---|
| Actinobacteria | 28 | 27.4 |
| Firmicutes | 28.6 | 26.4 |
| Proteobacteria | 23.6 | 24.9 |
| Acidobacteria | 6.7 | 7.4 |
| Chloroflexi | 5.5 | 6.8 |
| Gemmatimonadetes | 2.4 | 2 |
| Planctomycetes | 2.3 | 2.1 |
| Bacteroidetes | 1.1 | 0.9 |
| Verrucomicrobia | 0.9 | 0.8 |
| Nitrospirae | 0.6 | 1 |
| Candidatus Tectomicrobia | 0.1 | 0.2 |
| Cyanobacteria | <0.1 | 0.1 |
| Candidatus Saccharibacteria | <0.1 | <0.1 |
| Thermodesulfobacteria | <0.1 | <0.1 |
| Deinococcus-Thermus | <0.1 | <0.1 |
| Fusobacteria | <0.1 | <0.1 |
| Fibrobacteres | <0.1 | <0.1 |
| Chlamydiae | <0.1 | <0.1 |

**Table S3.** Relative abundance (%) in nodule microbiomes from four different Pakistan soils.

| Phylum | Genus | Family | sn1 | sn2 | sn3 | sn4 |
|---|---|---|---|---|---|---|
| Proteobacteria | *Bradyrhizobium* | *Bradyrhizobiaceae* | 86.2 | 78.9 | 85.7 | 88.8 |
| Proteobacteria | *Bosea* | *Bradyrhizobiaceae* | 3.2 | 3.1 | 3.3 | 2.9 |
| Proteobacteria | *Rhodopseudomonas* | *Bradyrhizobiaceae* | 1.8 | 1.5 | 1.6 | 1.7 |
| Proteobacteria | *Afipia* | *Bradyrhizobiaceae* | 0.9 | 1.0 | 0.8 | 0.8 |
| Proteobacteria | *Oligotropha* | *Bradyrhizobiaceae* | 0.2 | 0.2 | 0.1 | 0.1 |
| Proteobacteria | *Sinorhizobium/Ensifer* | *Rhizobiaceae* | 1.4 | 7.4 | 1.8 | 0.2 |
| Proteobacteria | *Rhizobium/Agrobacterium* | *Rhizobiaceae* | 0.2 | 0.8 | 0.2 | - |
| Proteobacteria | *Shinella* | *Rhizobiaceae* | 0.1 | 0.1 | 0.1 | 0.1 |
| Proteobacteria | *Rhodoplanes* | *Hyphomicrobiaceae* | 2.3 | 2.0 | 2.1 | 2.3 |
| Proteobacteria | *Mesorhizobium* | *Phyllobacteriaceae* | 0.1 | 0.4 | 0.2 | 0.2 |
| Proteobacteria | *Acidiphilium* | *Acetobacteraceae* | 0.2 | 0.2 | 0.2 | 0.2 |
| Proteobacteria | *Pseudomonas* | *Pseudomonadaceae* | 0.1 | 0.5 | 0.1 | 0.1 |
| Proteobacteria | Proteobacteria (others) | | 3.2 | 3.8 | 3.4 | 2.8 |
| | Others | | 0.1 | 0.2 | 0.2 | - |

**Table S4.** Morphological and physiological characterization of bacterial endophytes from cowpea nodule

| **Bacterial isolates** | **Media used*** | **Soil type/ plant genotype** | **Colony Morphology** | | | | **Gram-staining** | **Cell shape** | **Bacterial strain identification** |
|---|---|---|---|---|---|---|---|---|---|
| | | | **Color** | **Form** | **Elevation** | **Margin** | | | |
| **CPN1** | TY | SS1/603 | Cream | Irregular | Raised | Entire | -ve | Rods | *Mesorhizobium* |
| **CPN2** | TY | SS1/603 | Off-white | Irregular | Flat | Undulate | -ve | Rods | *Ensifer* |
| **CPN3** | TY | SS1/603 | Off-white | Irregular | Raised | Entire | +ve | Rods | *Paenibacillus* |
| **CPN4** | TSA | SS1/603 | Off-white | Circular | Umbonate | Entire | +ve | Rods | *Paenibacillus* |
| **CPN5** | TSA | SS2/603 | Off-white | Irregular | Raised | Entire | +ve | Rods | *Paenibacillus* |
| **CPN6** | TY | SS2/603 | Cream | Circular | Raised | Undulate | -ve | Rods | *Pseudomonas* |
| **CPN7** | TSA | SS2/603 | Cream | Circular | Raised | Entire | +ve | Filamentous | *Streptomyces* |
| **CPN8** | TSA | SS2/603 | Bright Yellow | Circular | Raised | Entire | +ve | Filamentous | *Streptomyces* |
| **CPN9** | TSA | SS3/603 | White | Circular | Flat | Entire | +ve | Filamentous | *Frankia* |
| **CPN10** | AG | SS3/603 | Off-white | Punctiform | Raised | Entire | -ve | Rods | *Bradyrhizobium* |
| **CPN11** | TY | SS3/603 | White | Irregular | Raised | Undulate | -ve | Rods | *Enterobacter* |
| **CPN12** | TY | SS3/603 | Off-white | Circular | Raised | Entire | -ve | Rods | *Sinorhizobium* |
| **CPN13** | TY | SS3/603 | Off-white | Irregular | Raised | Undulate | -ve | Rods | *Sinorhizobium* |

| | | | | | | | | | |
|---|---|---|---|---|---|---|---|---|---|
| **CPN14** | TSA | SS4/603 | Cream | Circular | Flat | Entire | -ve | Rods | *Pseudomonas* |
| **CPN15** | TSA | SS4/603 | Cream | Circular | Raised | Entire | +ve | Rods | *Bacillus* |
| **CPN16** | TY | SS4/603 | Off-white | Circular | Raised | Entire | -ve | Rods | *Pseudomonas* |
| **CPN17** | TSA | SS4/603 | Off-white | Irregular | Raised | Undulate | +ve | Rods | *Pseudomonas* |
| **CPN18** | TSA | SS1/603S | Light Yellow | Circular | Umbonate | Entire | -ve | Rods | *Pseudomonas* |
| **CPN19** | TY | SS1/603S | cream | Circular | Raised | Undulate | -ve | Rods | *Aeromonas* |
| **CPN20** | TY | SS1/603S | Off-white | Irregular | Raised | Entire | -ve | Rods | *Enterobacter* |
| **CPN21** | AG | SS1/603S | Light grey | Circular | Flat | Undulate | -ve | Rods | *Bradyrhizobium* |
| **CPN22** | TSA | SS1/603S | Off-white | Punctiform | Raised | Entire | +ve | Rods | *Bacillus* |
| **CPN23** | TY | SS2/603S | White | Irregular | Raised | Undulate | -ve | Rods | *Sinorhizobium* |
| **CPN24** | TSA | SS2/603S | Off-white | Circular | Raised | Entire | -ve | Rods | *Pseudomonas* |
| **CPN25** | TY | SS2/603S | Off-white | Irregular | Raised | Undulate | -ve | Rods | *Bradyrhizobium* |
| **CPN26** | AG | SS2/603S | Cream | Circular | Flat | Entire | -ve | Rods | *Bradyrhizobium* |
| **CPN27** | TY | SS3/603S | Cream | Circular | Raised | Entire | -ve | Rods | |
| **CPN28** | TY | SS3/603S | Off-white | Circular | Raised | Entire | -ve | Rods | *Bradyrhizobium* |
| **CPN29** | AG | SS3/603S | Off-white | Irregular | Raised | Undulate | +ve | Rods | *Bradyrhizobium* |

| | | | | | | | | | |
|---|---|---|---|---|---|---|---|---|---|
| **CPN30** | TSA | SS3/603S | Off-white | Irregular | Raised | Undulate | +ve | Rods | *Paenibacillus* |
| **CPN31** | TSA | SS4/603S | Off-white | Irregular | Raised | Undulate | +ve | Rods | *Paenibacillus* |
| **CPN32** | TY | SS4/603S | Off-white | Punctiform | Raised | Entire | -ve | Rods | *Pseudomonas* |
| **CPN33** | TSA | SS4/603S | White | Irregular | Raised | Undulate | -ve | Rods | *Enterobacter* |
| **CPN34** | TSA | SS4/603S | Off-white | Circular | Raised | Entire | +ve | Rods | *Bacillus* |
| **CPN35** | TSA | SS1/CB46S | Off-white | Irregular | Raised | Undulate | +ve | Rods | *Bacillus* |
| **CPN36** | TSA | SS1/CB46S | Cream | Circular | Flat | Entire | +ve | Rods | *Bacillus* |
| **CPN37** | TY | SS1/CB46S | Cream | Circular | Raised | Entire | +ve | Rods | *Paenibacillus* |
| **CPN38** | AG | SS1/CB46S | Off-white | Circular | Raised | Entire | +ve | Rods | *Paenibacillus* |
| **CPN39** | TY | SS2/CB46S | Off-white | Irregular | Raised | Undulate | -ve | Rods | *Mesorhizobium* |
| **CPN40** | TY | SS2/CB46S | Light Yellow | Circular | Umbonate | Entire | -ve | Rods | *Mesorhizobium* |
| **CPN41** | TSA | SS2/CB46S | Cream | Circular | Raised | Undulate | -ve | Rods | *Bradyrhizobium* |
| **CPN42** | TY | SS2/CB46S | Off-white | Irregular | Raised | Entire | -ve | Rods | *Bradyrhizobium* |
| **CPN43** | TY | SS3/CB46S | Light grey | Circular | Flat | Undulate | -ve | Rods | *Mesorhizobium* |
| **CPN44** | TSA | SS3/CB46S | Off-white | Circular | Raised | Entire | -ve | Rods | *Pseudomonas* |

| | | | | | | | | | |
|---|---|---|---|---|---|---|---|---|---|
| **CPN45** | TSA | SS3/CB46S | Off-white | Irregular | Raised | Undulate | -ve | Rods | *Mesorhizobium* |
| **CPN46** | TY | SS3/CB46S | Light Yellow | Circular | Umbonate | Entire | +ve | Rods | *Bacillus* |
| **CPN47** | AG | SS3/CB46S | Cream | Circular | Raised | Undulate | -ve | Rods | *Enterobacter* |
| **CPN48** | TSA | SS4/CB46S | Off-white | Irregular | Raised | Entire | +ve | Rods | *Bacillus* |
| **CPN49** | TY | SS4/CB46S | Cream | Circular | Flat | Undulate | -ve | Rods | *Pseudomonas* |
| **CPN50** | TSA | SS4/CB46S | Off-white | Circular | Umbonate | Entire | -ve | Rods | *Mesorhizobium* |
| **CPN51** | TSA | SS4/CB46S | Light Yellow | Circular | Raised | Undulate | +ve | Rods | *Bacillus* |

*TSA, Tryptone Soya Agar; TY, Tryptone Yeast extract Agar; AG, Arginine Glycerol Agar; SS1/603, Soil Sample1/cowpea 603; SS2/603, Soil Sample2/cowpea 603; SS3/603, Soil Sample3/cowpea 603; SS4/603, Soil Sample4/cowpea 603; SS1/603S, Soil Sample1/cowpea 603 under salinity stress; SS2/603S, Soil Sample2/cowpea 603 under salinity stress; SS3/603S, Soil Sample3/cowpea 603 under salinity stress; SS4/603S, Soil Sample4/cowpea 603 under salinity stress; SS1/CB46S, Soil Sample1/cowpea CB46 under salinity stress; SS2/CB46S, Soil Sample2/cowpea CB46 under salinity stress; SS3/CB46S, Soil Sample3/cowpea CB46 under salinity stress; SS4/CB46S, Soil Sample4/cowpea CB46 under salinity stress

**Table S5.** Antifungal activity of bacterial strains isolated from cowpea nodules.

| **Bacterial Isolates** | | **Antifungal activity** | | | | |
|---|---|---|---|---|---|---|
| | | ***Fusarium oxysporum*** | ***Fusarium solani*** | ***Curvularia* sp.** | ***Alternaria solani*** | ***Aspergillus flavus*** |
| **CPN1** | *Mesorhizobium* | - | - | - | - | - |
| **CPN2** | *Ensifer* | - | - | + | - | ++ |
| **CPN3** | *Paenibacillus* | +++ | + | ++ | +++ | - |
| **CPN4** | *Paenibacillus* | ++ | - | + | - | - |
| **CPN5** | *Paenibacillus* | - | - | - | ++ | ++ |
| **CPN6** | *Pseudomonas* | + | ++ | - | - | - |
| **CPN7** | *Streptomyces* | - | - | - | - | - |
| **CPN8** | *Streptomyces* | - | - | + | - | - |
| **CPN9** | *Frankia* | - | - | - | - | - |
| **CPN10** | *Bradyrhizobium* | - | + | ++ | - | + |
| **CPN11** | *Enterobacter* | ++ | - | - | - | - |
| **CPN12** | *Sinorhizobium* | - | - | ++ | - | - |
| **CPN13** | *Sinorhizobium* | - | - | - | - | - |
| **CPN14** | *Pseudomonas* | - | + | +++ | - | ++ |
| **CPN15** | *Bacillus* | ++ | + | - | ++ | - |
| **CPN16** | *Pseudomonas* | - | - | ++ | - | - |
| **CPN17** | *Pseudomonas* | - | + | - | - | - |
| **CPN18** | *Pseudomonas* | +++ | + | +++ | - | ++ |
| **CPN19** | *Aeromonas* | - | - | - | - | - |
| **CPN20** | *Klebsiella* | - | ++ | +++ | - | ++ |

| | | | | | | |
|---|---|---|---|---|---|---|
| **CPN21** | *Bradyrhizobium* | - | - | - | - | - |
| **CPN24** | *Bacillus* | - | - | - | - | - |
| **CPN27** | *Mesorhizobium* | - | - | - | - | - |
| **CPN30** | *Paenibacillus* | - | - | + | - | - |
| **CPN35** | *Bacillus* | +++ | ++ | +++ | - | ++ |
| **CPN36** | *Bacillus* | - | - | - | - | - |
| **CPN37** | *Paenibacillus* | - | + | - | - | - |
| **CPN38** | *Paenibacillus* | ++ | ++ | + | - | - |
| **CPN41** | *Bradyrhizobium* | - | - | - | - | - |
| **CPN43** | *Mesorhizobium* | + | - | ++ | - | +++ |
| **CPN44** | *Pseudomonas* | - | - | - | - | - |
| **CPN45** | *Mesorhizobium* | - | - | - | - | - |
| **CPN47** | *Bacillus* | ++ | + | - | ++ | - |
| **CPN48** | *Enterobacter* | - | - | - | - | - |

**Note: -, no activity; +, low activity; ++, medium activity; +++, high activity**

**Table S6:** Screening of hydrolytic enzymes produced by cowpea nodule associated bacterial strains

| Bacterial isolates | Hydrolytic enzymes activity | | | | |
|---|---|---|---|---|---|
| | Cellulase | Lipase | Chitinase | Amylase | Protease |
| CPN1 | + | - | - | + | ++ |
| CPN2 | - | - | - | + | - |
| CPN3 | +++ | - | ++ | +++ | +++ |
| CPN4 | + | - | + | - | ++ |
| CPN5 | + | - | ++ | + | - |
| CPN6 | - | ++ | - | - | - |
| CPN7 | - | + | - | + | ++ |
| CPN8 | ++ | - | + | ++ | ++ |
| CPN9 | - | - | ++ | - | ++ |
| CPN10 | - | - | ++ | ++ | - |
| CPN11 | ++ | - | + | ++ | + |
| CPN12 | ++ | - | - | + | ++ |
| CPN13 | + | - | - | + | ++ |
| CPN14 | - | ++ | +++ | ++ | + |
| CPN15 | + | - | ++ | + | ++ |
| CPN16 | - | ++ | - | ++ | - |
| CPN17 | ++ | - | + | - | - |
| CPN18 | ++ | +++ | +++ | - | +++ |
| CPN19 | +++ | + | - | ++ | - |
| CPN20 | +++ | - | +++ | ++ | ++ |
| CPN21 | + | - | - | ++ | + |
| CPN24 | - | ++ | - | - | + |

| | | | | | |
|---|---|---|---|---|---|
| **CPN27** | + | - | - | ++ | ++ |
| **CPN30** | - | - | ++ | ++ | + |
| **CPN35** | +++ | - | +++ | - | ++ |
| **CPN36** | ++ | + | - | - | ++ |
| **CPN37** | + | - | ++ | - | - |
| **CPN38** | ++ | - | - | ++ | - |
| **CPN41** | - | - | ++ | ++ | - |
| **CPN43** | - | + | + | ++ | - |
| **CPN44** | ++ | - | ++ | ++ | - |
| **CPN45** | + | - | - | - | + |
| **CPN47** | ++ | - | ++ | + | ++ |
| **CPN48** | ++ | - | +++ | - | + |

**Note: -, no activity; +, low activity; ++, medium activity; +++, high activity**
